# The cortico-striatal circuit regulates sensorimotor gating via Disc1/Huntingtin-mediated Bdnf transport

**DOI:** 10.1101/497446

**Authors:** Hanna Jaaro-Peled, Sunil Kumar, Dalton Hughes, Sun-Hong Kim, Sandra Zoubovsky, Yuki Hirota-Tsuyada, Diana Zala, Akiko Sumitomo, Julie Bruyere, Brittany M. Katz, Beverly Huang, Rafael Flores, Soumya Narayan, Zhipeng Hou, Aris N. Economides, Takatoshi Hikida, William C. Wetsel, Karl Deisseroth, Susumu Mori, Nicholas J. Brandon, Motomasa Tanaka, Koko Ishizuka, Miles D. Houslay, Frédéric Saudou, Kafui Dzirasa, Akira Sawa, Toshifumi Tomoda

**Author notes:** To whom correspondence should be addressed: Akira Sawa,; Kafui Dzirasa,; Toshifumi Tomoda. These authors contributed equally: Hanna Jaaro-Peled and Sunil Kumar.

## Abstract

Sensorimotor information processing that underlies normal cognitive and behavioral traits is dysregulated across a subset of neurological and psychiatric disorders. The cross-disease deficit in sensorimotor gating poses a unique opportunity to integrate hierarchical findings at molecular, cellular, through circuitry levels to obtain an in-depth mechanistic understanding of this process that contributes to brain physiology and pathophysiology beyond categorical segmentation of brain disorders. Based on circuitry recording with wild-type mice, we demonstrated that the cortico-striatal projection mediates sensorimotor gating responses during prepulse inhibition (PPI) task. We also found that these circuitry responses were disrupted in *Disc1* locus-impairment (LI) mice, a model representing neuropsychiatric conditions. Thus, we hypothesized that Disc1-mediated molecular and cellular machinery along the cortico-striatal circuit may regulate sensorimotor gating. Anatomical and biochemical analyses of *Disc1-*LI mice revealed attenuated Bdnf transport along the cortico-striatal circuit. Pharmacologically augmenting Bdnf transport by chronic lithium administration, in part via Ser-421 phosphorylation of Huntingtin (Htt) and its integration into the motor machinery, restored the striatal Bdnf levels and PPI deficits in *Disc1*-LI mice, suggesting that the Bdnf transport attenuation mechanistically underlies the circuitry and behavioral deficits. These results also shed light on a novel mechanism and utility of lithium that is currently used as a major mood stabilizer in clinical settings. Collectively, the present study illustrates integrative biological mechanisms for sensorimotor gating, underscoring the cross-disease nature of this behavioral dimension and translational utility of the findings under the era of precision medicine in brain disorders.

## Introduction

To address a fundamental biological question of how the brain drives behaviors under physiological and pathophysiological conditions, a wide range of efforts have been historically utilized. One popular approach is to focus on a specific neuropsychiatric disorder and analyze functional outcomes by perturbing a critical genetic factor or biological mediator for the disease. This approach is proven scientifically effective when the disease diagnosis is defined by biological and etiological evidence, e.g., in the case of Huntington’s disease caused by genetic alterations in *Huntingtin (HTT)* gene. However, in reality, the diagnostic criteria for most psychiatric disorders are formally defined to achieve clinical reliability by sacrificing etiological validity (1). As a result, biologically heterogeneous conditions are included in each diagnosis in this “categorical” approach. To overcome this limitation, recent discussion in the clinical nosology of brain disorders has brought a “dimensional” approach, in which the mechanism for a critical behavioral trait or dimension, independent of a diagnostic category, is addressed at multiple levels (e.g., molecular, cellular, and circuitry levels) (2). In this approach, behavioral dimensions that are directly translatable between humans and model animals, such as rodents, are particularly appreciated from both basic and clinical neuroscience viewpoints.

Sensorimotor gating is one of these representative dimensions; it has classically been measured using prepulse inhibition (PPI) of a startle response. During the PPI test, neurophysiological responses obtained following presentation of a startle stimulus are compared to the responses obtained to the same startle stimulus when it is preceded by a lower amplitude prepulse stimulus (3). In normal individuals, presentation of a prepulse stimulus suppresses both the normal neurophysiological and behavioral responses to the startle stimulus. Conversely, PPI is diminished in individuals with a wide range of neuropsychiatric disorders (4). Furthermore, PPI is conserved across multiple different species (5) and deficits in PPI are observed in a series of genetic- and pharmacological-based animal models that are developed to understand mechanisms for psychosis or mood dysregulation (6-8). Multiple circuits involving the forebrain are hypothesized to modulate the inhibitory functions of the prepulse (4). Preclinical studies have demonstrated that striatal lesions disrupt PPI (9). Other studies have suggested the involvement of the prefrontal cortex (PFC) in PPI (10). Nevertheless, as far as we are aware, studies that directly address the role of cortico-striatal circuitry for PPI are unavailable. It remains elusive how PPI and sensorimotor gating are mechanistically explained as an integrative behavioral dimension at the circuitry, and down to the cellular and molecular levels. Given its deficits reported across brain dysfunctions, such as schizophrenia, psychotic disorders, mood disorder, and neurodegenerative disorders, elucidating the mechanisms of PPI will have high clinical and social impacts.

In the present study, our aim is to comprehensively understand a key mechanism for PPI and sensorimotor gating at molecular, cellular, circuit, and behavioral levels by taking a hypothesis-driven dimensional approach in an integrative manner. By using microwire arrays chronically implanted in mouse brains combined with optogenetics, we functionally show for the first time that the cortico-striatal projection mediates the PPI. We next explore molecular leads that underlie the circuitry mechanism. At the molecular level, multiple groups, including our team, have previously reported that PPI deficits are associated with perturbation of Disc1 (11-13) and Huntingtin (Htt) (14), which interact with each other at the protein level (15). Htt is the causal factor for Huntington’s disease, and the patients show PPI deficits (16). Likewise, a balanced translocation that disrupts the *DISC1* gene is reportedly linked to a wide range of major mental illnesses, such as schizophrenia and depression, in a Scottish pedigree (17). Thus, we hypothesized that DISC1 protein, interacting with Htt protein, may regulate the cortico-striatal circuitry mechanism for the sensorimotor gating responses that underlie a wide range of neuropsychiatric conditions (18). Here we first report electrophysiological disturbances of the cortico-striatal circuitry, which mediates the PPI, in a *Disc1* genetic model. By studying this model further, we now demonstrate how the protein interaction of Disc1 and uniquely phosphorylated Htt at serine-421 contributes to proper function of this cortico-striatal circuitry via Bdnf transport, which is required for sensorimotor gating responses at the cellular, circuitry and behavioral levels during the PPI task.

## Results

### Critical roles for the PFC and DMS in sensorimotor gating: studies with wild-type mice

Previous studies reported that PPI was affected by manipulation of the PFC or the DMS (9, 10). Thus, we hypothesized that the cortico-striatal projection might mediate sensorimotor gating. To test this hypothesis, wild-type (WT) C57BL/6 mice were implanted with microwire arrays, allowing us to simultaneously record single unit activities as well as local field potentials (LFPs) from the PFC and DMS in awake, partially-restrained conditions *(i.e*., in a plexi-glass tube) during PPI testing, in which startle trial (120 dB) and prepulse stimulus followed by startle stimulus (PPI trial) were randomly given to each mouse (**Figure 1A**). All implantation sites were verified histologically after completing the *in vivo* recording (**Figure S1**).

**Fig. 1.**
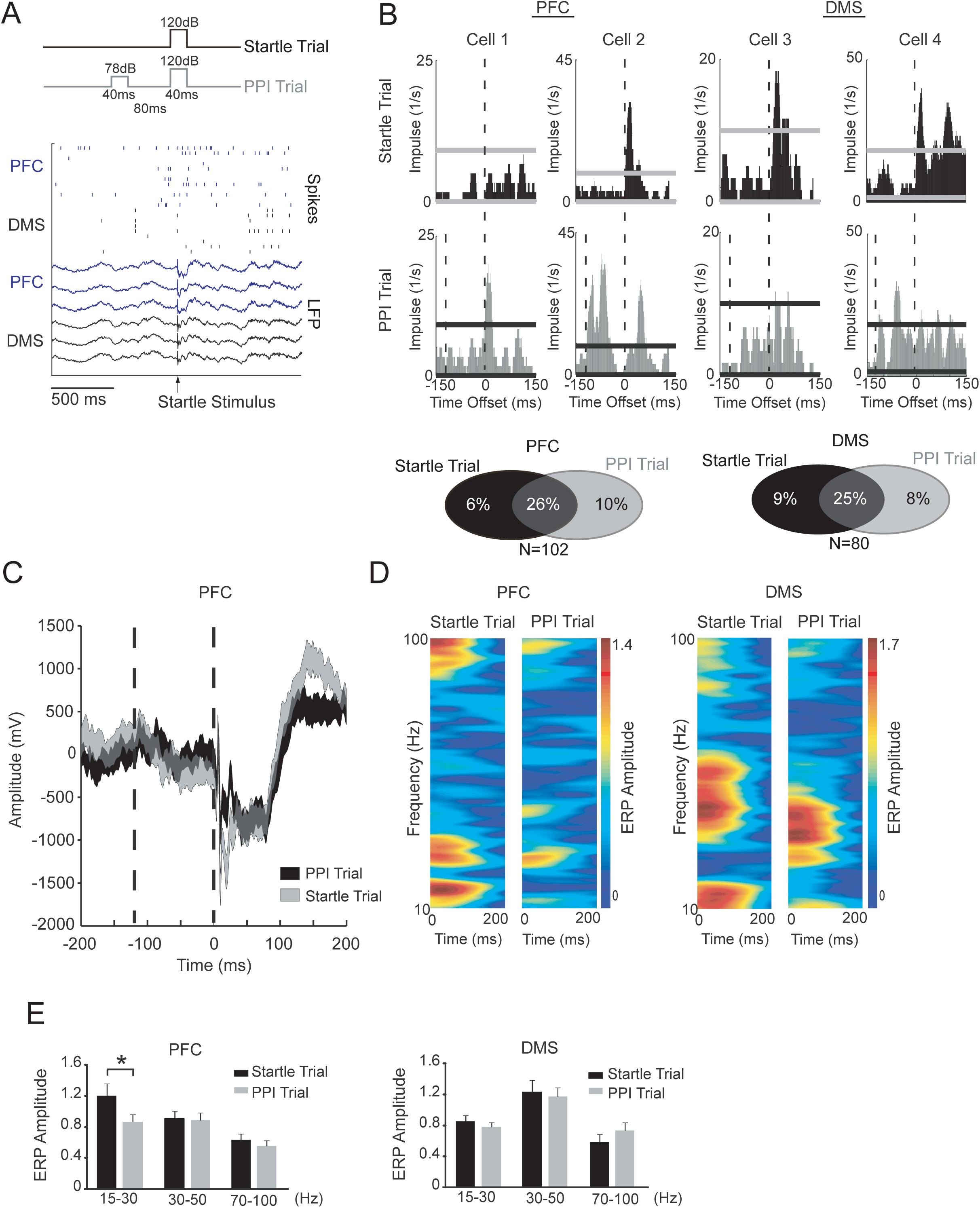
The prefrontal cortex and dorsomedial striatum in sensorimotor gating. (**A**) Startle (120 dB) and pre-pulse (78 dB) stimuli used for the PPI task (top). Two second trace of single neuron spikes and LFP (local field potential) activity recorded in a mouse during the PPI test (bottom). (**B**) Perievent time histograms showing examples of PFC (left) and DMS (right) unit responses during startle and PPI trials (n = 30 per trial, data are shown in 20ms bins). Dashed lines correspond to the presentation of the 78dB stimulus (−120ms) and the 120dB stimulus (0 ms). The percentage of neurons that responded to the startle stimulus during the startle and PPI trials are shown below. (C) PFC local field potentials recorded during prepulse and startle trials. Zero milliseconds (ms) corresponds to the time of presentation of the 120dB pulse for both trial types. Data are shown as mean ± SEM (n = 30 trials for each stimulus). (D) Amplitude-frequency components of PFC (left) and DMS (right) local field potentials (LFP) normalized to the mean LFP amplitudes observed during the −5s to −1s window prior to the presentation of the 120 dB pulse. (E) The prepulse significantly reduced the mean cortical (left), but not the DMS (right) beta response to the startle pulse (data were averaged within animals across 8-16 LFP channels per brain area); *P<0.05 using mixed-model ANOVA followed by Bonferonni-corrected Wilcoxon sign-rank test.

In the single unit activity recording, we observed that a significant proportion of neurons responded to the startle stimulus of the startle trials (32% of 102 PFC cells and 34% of 80 DMS cells) and a significant portion of neurons responded to the startle stimulus during the PPI trials (36% of 102 PFC cells and 33% of 80 DMS cells) (**Figure 1B**). While 26% of PFC and 25% of DMS cells responded to both trial types, 16% of PFC neurons and 17% of DMS neurons responded to the startle stimulus exclusively during one trial type (startle trial or PPI trial), but not the other. Thus, both PFC and DMS activities reflected a neurophysiological correlate of sensory gating processes taking place during the PPI test. We next quantified the effect of the gating stimulus (78 dB, low amplitude prepulse stimulus) on LFP responses to the startle stimulus. Specifically, we calculated the mean evoked potential for each trial type and normalized the mean amplitude for each frequency to the amplitude observed during the 1-5 second interval prior to delivering the startle pulse (**Figure 1C, D**). This frequency-wise analysis approach allowed us to quantify the impact of the prepulse stimulus on the oscillatory activity induced by the startle stimulus. Using this strategy, we found that the prepulse stimulus significantly reduced the induction of PFC beta oscillations (15-30 Hz) elicited by the initial startle stimulus (**Figure 1E**). Taken together with our single cell observations, these analyses provided evidence that both the cortex and striatum signal the sensorimotor gating response.

### Critical role of the PFC-DMS projection in sensorimotor gating: comparative studies with wild-type and *Disc1* LI mice

Based on the results above, we next hypothesized that the cortico-striatal pathway directly regulates PPI and sensorimotor gating. To address this question, we compared WT mice with a genetically-engineered model that displays deficits in the sensorimotor gating response. In the present study, we chose a model in which *Disc1* is perturbed. Robust deficits in the PPI test for sensorimotor gating were indeed confirmed in a model with a deletion at the *Disc1* locus (Disc1 LI model, −/−) (**Figure 2A**) (19, 20). Structural magnetic resonance imaging (sMRI), a technique that addresses gross anatomical changes in an unbiased manner, showed a significant volume reduction only in the striatum and cerebellum among multiple brain regions in *Disc1* LI mice compared with controls (**Figure S2**). In analogy to the modest but reproducible changes in brain anatomy found in patients with mental illnesses (21), the volume reduction in the striatum of *Disc1* LI mice is mild, but significantly different compared with littermate controls (**Figure S2**).

**Figure 2.**
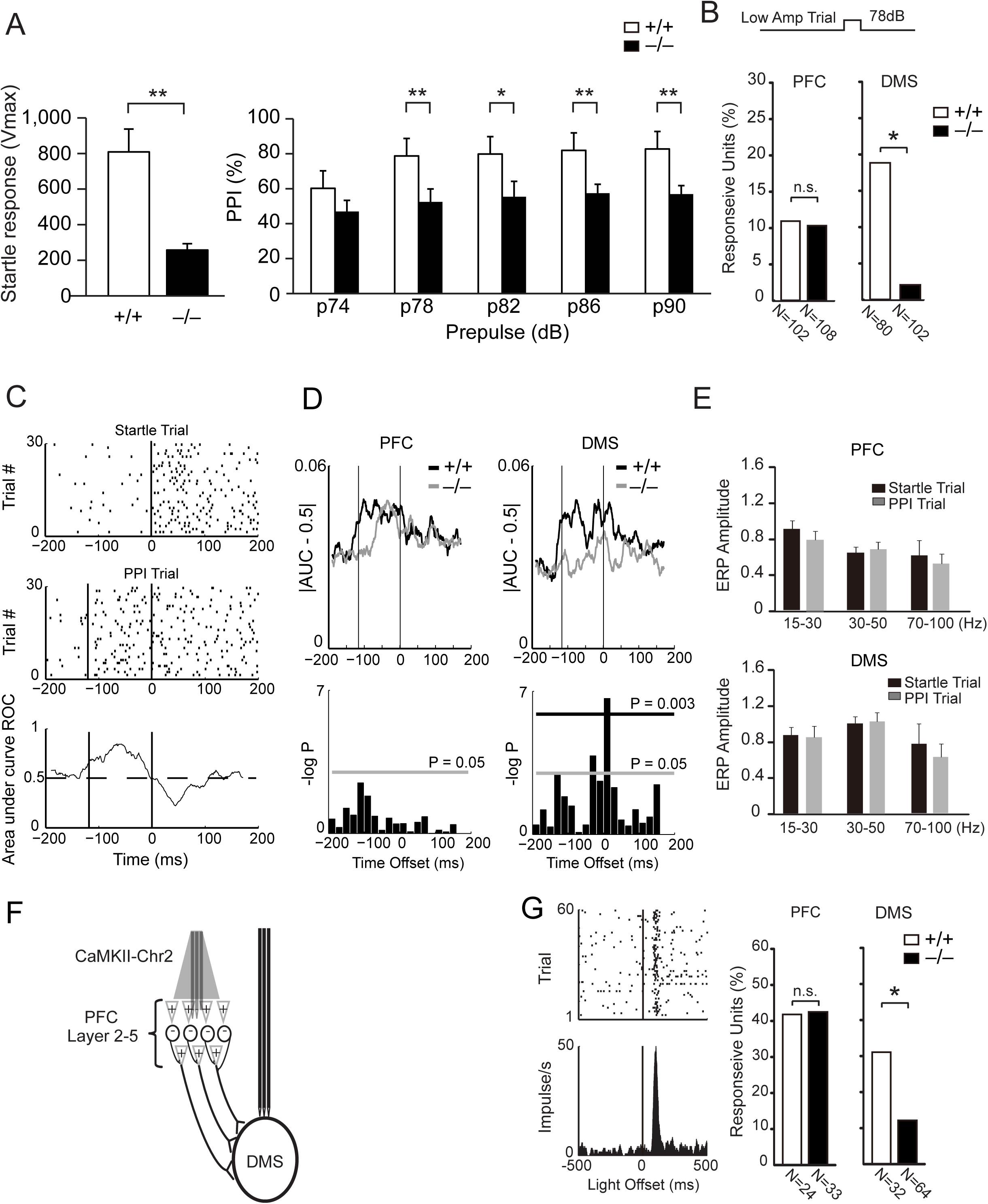
Sensorimotor gating in *Disc1* LI mice. (A) *Disc1* LI (-/-) mice show reduced startle response (left) and PPI (right). WT, n = 7; *Disc1* LI, n = 8. Data are shown as mean ±SEM. **p<0.01; *p<0.05 using student’s t-test. (B) Startle and prepulse stimuli used for the PPI task (top). A similar portion of PFC neurons modulated their firing rates in response to the 78dB low amplitude stimulus in *Disc1* LI (−/−) mice and their WT (+/+) littermates (P>0.05 using Z-test). A significantly lower proportion of DMS neurons modulated their response to this stimulus in *Disc1* LI mice compared to their WT littermates (*P<0.05 using Z-test). (C) Raster plot of DMS neuron response during startle and prepulse trials (top). Area under the ROC curve (AUC) demonstrating unit detection of gating across the stimulus interval (bottom). (D) Population AUC magnitude functions in WT and *Disc1* LI mice (top). Differences between genotypes were identified by comparing AUC functions averaged within 20ms bins using a Wilcoxon rank-sum test (bottom). The gray line corresponds with P=0.05. The black line corresponds with the significance threshold following Bonferroni correction for multiple comparisons (N=102 PFC neurons and 80 DMS neurons in WT mice. N=108 PFC neurons and 102 DMS neurons in mutant mice). (E) The prepulse stimulus showed no effect on the mean PFC (top) or DMS (bottom) response to the startle stimulus across any frequency band examined (data was averaged within animal across 8-16 LFP channels per brain area, n = 10 mice). (F) Schematic of concurrent optogenetic stimulation and neurophysiological recordings in *Disc1* LI and WT mice infected with AAV-CaMKII-Chr2 in PFC. (G) Sixty light pulses (10ms pulse width) were delivered with a pseudorandomized inter-pulse-interval ranging between 8 and 23 s. Left: Raster plot (top) and firing rate perievent time histogram (PETH) of a representative striatal neuron (bottom). Right: A similar proportion of PFC neurons modulated their firing rates in response to cortical stimulation (P>0.05 using Z-test) in *Disc1* LI (−/−) mice and WT (+/+) littermates. A significantly lower proportion of DMS neurons modulated their firing rates in response to cortical stimulation (right; *P<0.05 using Z-test) in *Disc1* LI mice compared to WT littermates.

Next, we tested the functional role of the cortico-striatal projection (or PFC-DMS projection) in PPI in *Disc1* LI mice compared with WT mice. To quantitatively address the function, we implanted them with microwire arrays and used the neurophysiological parameters of the cortical and striatal activities that were used in the analysis of WT mice (**Figure 1**). When a low amplitude acoustic stimulus (78 dB) was provided during the PPI testing sequence, a similar percentage of PFC neurons were activated in both *Disc1* LI and the WT mice (11/102 and 11/108 cells for WT and *Disc1* LI mice, respectively), whereas DMS activation following auditory stimulation was diminished in *Disc1* LI compared with WT mice (15/80 and 2/102 cells for WT and *Disc1* LI mice, respectively; **Figure 2B**). To further verify differences in neuronal sensory gating signals, we used a measurement based on receiver operator characteristic (ROC) analysis, which is widely used for classifying neurons’ responses to stimuli (22). In contrast to the analysis applied to averages of populations of neurons (**Figure 2B**), ROC analysis provides an estimate of how the activity of a single neuron differs between two stimuli on a trial-by-trial basis (i.e. the quality of signal detection; **Figure 2C**). Using this analysis, we found that *Disc1* LI mice showed diminished striatal signaling of sensory gating compared to WT littermates (**Figure 2D**). Cortical sensory gating was lower in the mutants as well, though these differences did not reach statistical significance (**Figure 2D**). Taken together with our results obtained from population analysis data, the data demonstrate functional deficits in the cortico-striatal circuit of *Disc1* LI mice, which may be causally related to the sensory gating deficits. We next quantified the effect of the gating stimulus (78 dB, low amplitude prepulse stimulus) on LFP responses to the startle stimulus in *Disc1* LI mice (the same way we did for WT mice in **Figure 1 C-E**). In contrast to the gating effect seen on PFC beta oscillations in WT mice (**Figure 1E**), the LFP responses in *Disc1* LI showed no differences between the startle trial and the PPI trial (**Figure 2E**), i.e. impaired gating.

To more directly test our hypothesis that *Disc1* LI mice may have deficits in the PFC-DMS cortico-striatal projection, we probed the projection with concurrent optogenetic and *in vivo* recording approaches (**Figure 2F**). We infected adeno-associated virus (AAV) encoding CaMKII-ChR2 into the PFC of *Disc1* LI mice or WT littermates, and then quantified neuronal activity of the PFC and DMS in response to light stimulation to the PFC (**Figure S3**). When light stimulation was delivered, a similar percentage of PFC neurons responded in both *Disc1* LI and WT mice (**Figure 2G**), demonstrating that there is no difference in the activation of the soma of PFC neurons between genotypes. On the other hand, striatal activation in response to cortical stimulation by light was significantly diminished in *Disc1* LI mice (**Figure 2G**). This shows that *Disc1* LI mice exhibit impaired function of the cortico-striatal projection in response to direct stimulation. Taken together, the electrophysiology experiments show that the cortico-striatal projection is critical for sensorimotor gating and that *Disc1* LI mice have deficient cortico-striatal gating.

### Role for striatal Bdnf through the PFC-DMS projection in sensorimotor gating

The striatum critically depends on a supply of Bdnf via its transport along projections from cortical neurons (23). Deficits in the supply of Bdnf to the striatum lead to a reduction in striatal volume (24). Thus, the selective reduction in striatal volume and functional deficits in cortical-striatal circuitry prompted us to test the amount of Bdnf in the striatum of *Disc1* LI mice. Therefore, we measured Bdnf by enzyme-linked immunosorbent assay (ELISA) in 1.5-, 3-, and 6-month-old mice. While no significant differences in cortical Bdnf levels were observed between *Disc1* LI and WT at any age tested, the levels of striatal Bdnf were significantly lower in *Disc1* LI mice starting from 3 months of age (**Figure 3A**). Notably, the cortical and striatal Bdnf mRNA levels in *Disc1* LI were equivalent to those in WT (**Figure 3B**). Thus, the data suggest a possibility that the decrease in striatal Bdnf may, at least in part, be due to deficits in Bdnf transport from the cortex to the striatum in *Disc1* LI mice. To further examine this possibility, we analyzed Bdnf transport in primary cortical neurons from *Disc1* LI and WT mice. The velocity of anterograde and retrograde Bdnf transport was reduced in *Disc1* LI compared with WT and was rescued by transfection of full-length Disc1 (**Figure 3C**).

**Figure 3.**
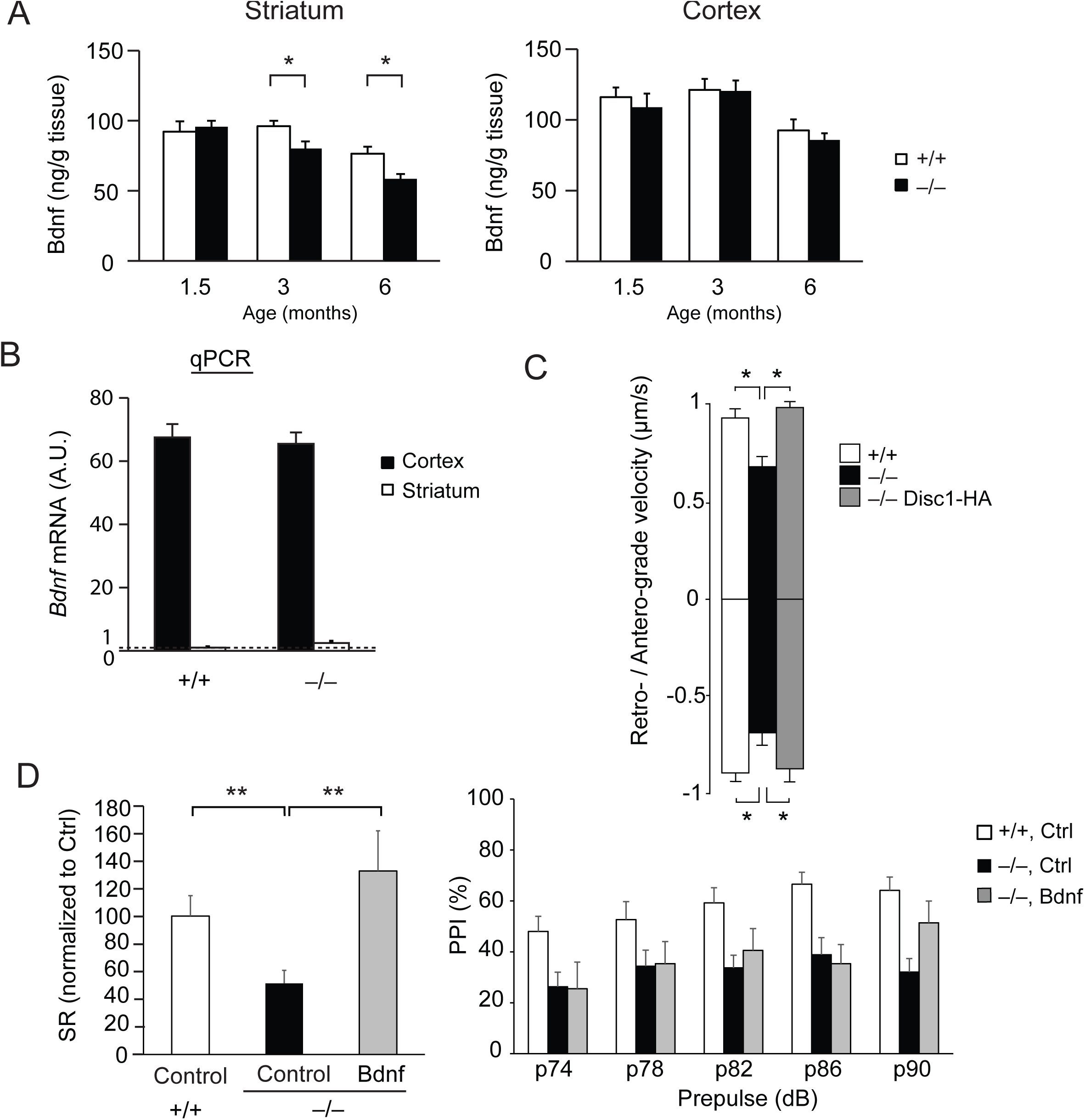
Deficits in corticostriatal trafficking of Bdnf in *Disc1* LI mice. (A) Age-dependent reduction in striatal, but not cortical, Bdnf in *Disc1* LI (−/−) mice as measured by ELISA. WT, n = 9-10; *Disc1* LI, n = 9-10. *P<0.05 using student’s t-test. (B) No difference in *Bdnf* mRNA in the cortex and striatum between WT (+/+) and *Disc1* LI (−/−) mice at 3 months of age. *Bdnf* mRNA in WT striatum was assigned as “1” to which the other results were normalized. WT, n = 4; *Disc1* LI, n = 4. (C) Impaired retrograde and anterograde trafficking velocity in primary cortical neurons, which could be rescued by over-expression of full-length Disc1 (−/− Disc1-HA). *P<0.05 using Kruskal-Wallis tests with Dunn’s multiple comparisons. (D) Left: AAV-Bdnf injection into the striatum normalized the startle response during the PPI task, compared with mock AAV injection (Control) in *Disc1* LI (−/−) mice. P < 0.01 using student’s t-test. Right: AAV-Bdnf tended to improve PPI with a prepulse at 90 dB (P = 0.06 using t-test). WT+control, n = 11; *Disc1* LI+control, n = 10-14; *Disc1* LI+Bdnf, n = 5-9. Data in all panels are shown as means ± SEM.

Our findings showing that *Disc1* LI mice have low striatal Bdnf and functional striatal deficits, as well as a recent report showing that heterozygous Bdnf mutant mice have attenuated PPI (25), suggest a causal link between low levels of Bdnf and the PPI deficit in *Disc1* LI mice. To address the extent to which the striatal Bdnf reduction underlies PPI deficits in this model, we supplemented Bdnf by bilateral stereotaxic injection of AAV-Bdnf to the striatum. Six weeks after the injection, when exogenous Bdnf was fully expressed (26), expression of Bdnf, but not the control AAV, normalized the startle response of *Disc1* LI mice to levels equivalent to those of WT, and tended to improve the PPI deficits during the prepulse trials at 90 dB (**Figure 3D**). These data suggest that a reduced Bdnf supply from the cortex to the striatum, at least in part, underlies PPI deficits in *Disc1* LI mice.

We previously reported that Disc1 plays a role in axonal transport of synaptic vesicles in culture (27), supporting the idea that Kaibuchi and colleagues originally proposed (28). We further demonstrated that lithium could ameliorate transport deficits elicited by *Disc1* knockdown in culture (27). Lithium is a representative mood stabilizer (29-31), and is reportedly neurotrophic and neuroprotective possibly through BDNF augmentation (32-34). We therefore tested whether chronic lithium treatment would normalize the PPI deficits and biochemical abnormalities in *Disc1* LI mice. Lithium (100 mg/kg, i.p., daily, 14 days) normalized the PPI deficits (**Figure 4A**), enhanced Bdnf transport in neurons from *Disc1* LI mice *in vitro* (**Figure 4B**), and increased the levels of striatal Bdnf in *Disc1* LI mice *in vivo* (**Figure 4C**). Collectively, the data suggest that the cortico-striatal Bdnf transport machinery, sensitive to lithium-mediated regulation, at least in part underlie the PPI-associated cortico-striatal circuitry function.

**Figure 4.**
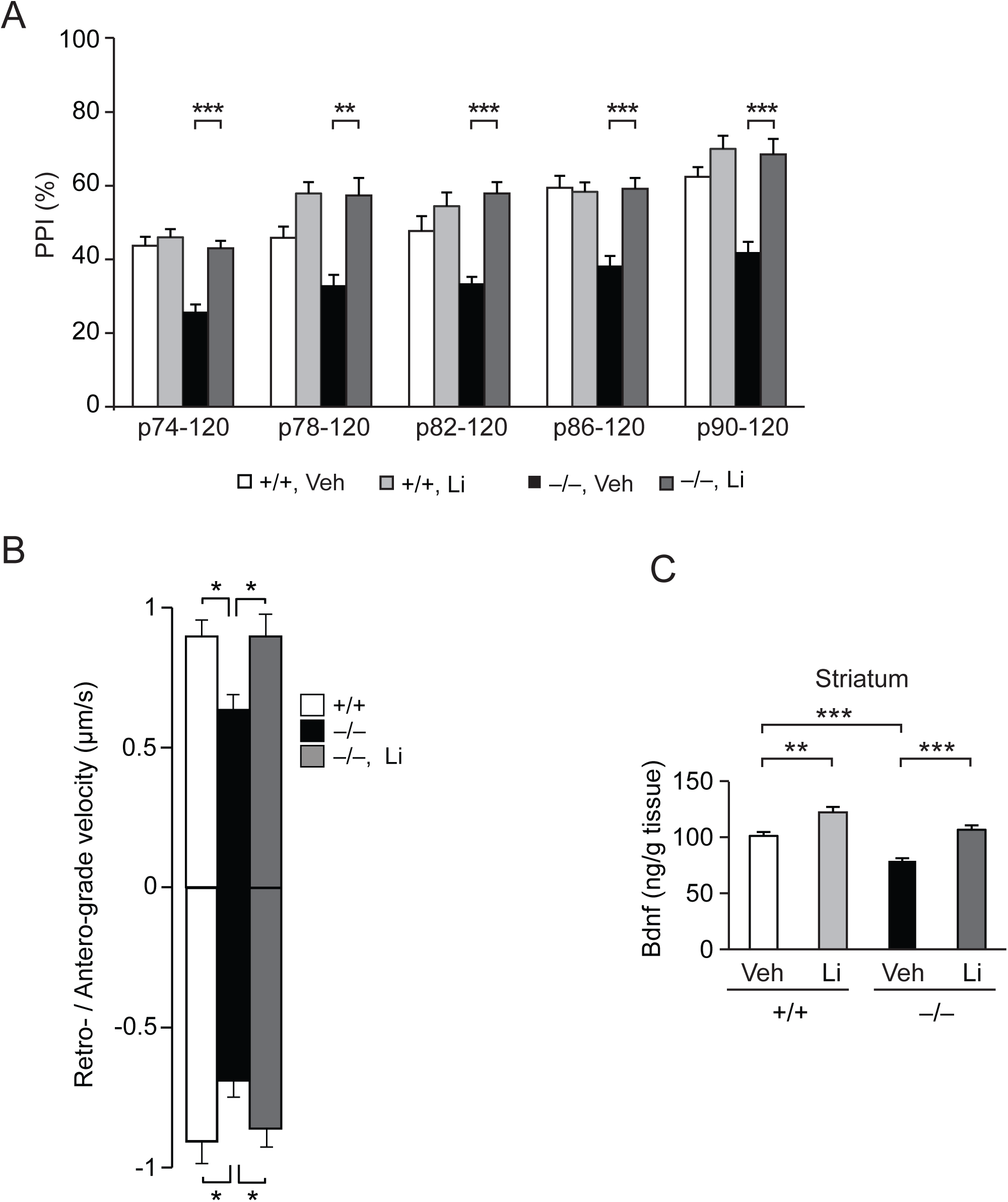
Lithium-mediated augmentation of Bdnf transport can rescue PPI deficits in *Disc1* LI mice. (A) Lithium (Li, 100 mg/kg body weight, i.p., daily, 14 days) normalized the PPI deficits in *Disc1* LI (−/−) mice. n = 5-8. Veh, vehicle; **P < 0.01, ***P< 0.001 using repeated measures two-way ANOVA with Bonferoni post-hoc test. (B) Li (2 mM in the culture media 30 min before imaging) normalized the slow retrograde/anterograde Bdnf transport in cultured primary neurons prepared from *Disc1* LI (−/−) mice. *P < 0.05 using Kruskal-Wallis tests with Dunn’s multiple comparisons. (C) Li (100 mg/kg body weight, i.p., daily, 14 days) increased the levels of Bdnf in the striatum of *Disc1* LI (−/−) mice to levels equivalent to WT (+/+) mice. The injections also increased Bdnf in WT, but the effects were more prominent in *Disc1* LI mice. **P < 0.01, ***P < 0.001.

### Disturbance of phospho-Huntingtin at serine-421: a mechanism for the deficit in the PFC-DMS transport of Bdnf and sensorimotor gating in *Disc1* mice

We have recently reported that Disc1 interacts with Huntingtin (Htt), the causal factor for Huntington’s disease (HD) at the protein level (15). Htt is a multifunctional protein, but one of its crucial roles is to regulate BDNF transport (35): phosphorylation of serine (Ser)-421 on Htt by Akt1 kinase is crucial for efficient BDNF transport (36, 37). In addition, lithium is reported to activate Akt1 in neurons (30). Therefore, we investigated a possible mechanism by which lithium augments Bdnf transport through Htt phosphorylation in *Disc1* LI mice.

We first confirmed that lithium (Li, 2 mM in the culture media, 16 h) activated Akt1 and significantly upregulated the phosphorylation of Htt Ser-421 in primary neurons in culture (**Figure 5A**). Importantly, we observed attenuated phosphorylation of Htt Ser-421 in the cortices of *Disc1* LI mice compared with WT (**Figure 5B**). Chronic lithium administration significantly upregulated the levels of Htt Ser-421 phosphorylation in *Disc1* LI cortices (**Figure 5C**). These results suggest that phosphorylation of Htt Ser-421 may be involved in the deficits of Bdnf transport in *Disc1* LI mice.

**Figure 5.**
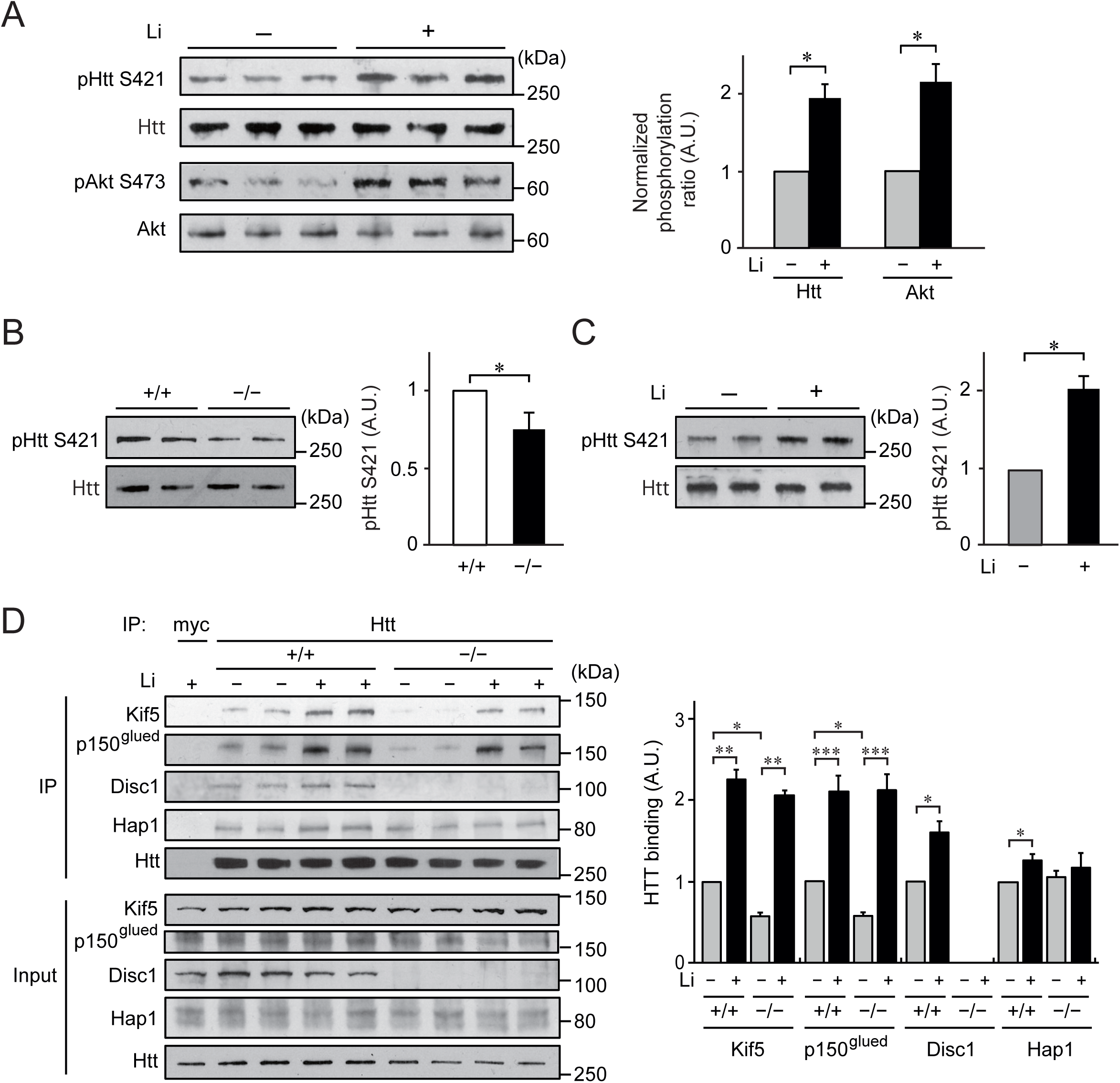
Lithium upregulates Htt Ser-421 phosphorylation and enhances the assembly of the Bdnf transport machinery in *Disc1* LI mice. (A) Lithium (Li, 2 mM in the culture media, 16 h) upregulated levels of phospho-Htt Ser-421, and phospho-Akt1 Ser-473 in primary cortical neurons. Levels of phosphoproteins were normalized by total levels of each protein. *p < 0.05 (Student’s t-test). (B) Relative phospho-Htt Ser-421 levels in *Disc1* LI (−/−) cerebral cortex at 3 months of age, normalized to those in WT (+/+) brains. *p < 0.05 (Student’s t-test). (C) Relative phospho-Htt Ser-421 levels in *Disc1* LI cerebral cortex at 3 months of age, normalized by total Htt levels, in the presence (+) or absence (−) of Li administration (100 mg/kg body weight, i.p., daily, 14 days). *P < 0.05 (Student’s t-test). (D) Cerebral cortical homogenates from WT (+/+) and *Disc1* LI (−/−) mice, chronically treated with (+) or without (−) Li (100 mg/kg body weight, i.p., daily, 14 days), were immunoprecipitated (IP) with anti-Htt antibody and analyzed by Western blots using the antibodies indicated. Graph: Relative binding capacity between Htt and each component of the Bdnf transport machinery. Each Western blot was done in triplicate. Data are shown as means ± SEM. *P < 0.05, **P < 0.01, and ***P < 0.001 (Student’s t-test).

To address the mechanism more precisely, we next evaluated additional molecular components of the Bdnf transport machinery in *Disc1* LI cortices in the presence or absence of lithium. Co-immunoprecipitation assays confirmed that some components of the Bdnf transport machinery [*i.e*., Kinesin heavy chain (KIF5), dynactin subunit p150^Glued^] (35) are less tightly assembled with Htt in *Disc1* LI brains compared with brains of normal controls (**Figure 5D**). Of importance, lithium treatment enhanced and normalized the interaction among these components (**Figure 5D**). Taken together, *Disc1* LI (or possibly depletion of key isoforms of the Disc1 protein) attenuates phosphorylation of Ser-421 on Htt and impairs the integration of Htt into the motor machinery, which negatively impacts Bdnf transport. Disc1 likely accounts for this integration mechanism, as Disc1 interacts with Htt and is included in the motor machinery responsible for Bdnf transport (**Figure 5D**). This mechanism, as schematically illustrated in **Figure S4**, is consistent with the idea that Ser-421 on Htt is phosphorylated by Akt1, a molecular target of lithium (30, 36, 37). In this model, both Disc1 and lithium can serve to facilitate the phosphorylation of Htt Ser-421 either by increasing the Akt1 accessibility to the motor complex or by upregulating Akt1 activity, which leads to the formation of a more stable motor complex responsible for Bdnf transport.

Note that, like other animal models, the behavioral deficits of *Disc1*-LI mice are not limited to those in the sensorimotor gating dimension (see **Figure S5**). They are hypoactive in the open field and impaired in the rotarod test, similar to many HD mouse models (38). As far as we are aware, however, none of the behavioral deficits in the other dimensions are likely to affect the PPI deficits. Thus, we believe that the present approach of focusing on one behavioral dimension with the *Disc1*-LI model is valid in that it could fully integrate underlying mechanisms from molecules to cellular, and circuitry mechanisms pertinent to a specific behavioral outcome.

## Discussion

Here we report a key mechanism underlying PPI and sensorimotor gating at the molecular, cellular, circuitry, and behavioral levels in an integrated manner by using a hypothesis-driven dimensional approach. We defined that the cortico-striatal (*i.e*., the PFC→DMS) projection is critical for normal PPI at the circuitry level and that the Disc1-containing motor complex, including Htt and Bdnf, in the PFC→DMS projection accounts for a key mechanism at the molecular level. Amelioration of the PPI deficits in the *Disc1*-LI model by lithium treatment is explained by augmentation of Bdnf transport via its pharmacological action on Akt1 activity and a specific phosphorylation of Htt at serine-421.

In addition to the contribution of this study to basic molecular and behavioral neuroscience, we believe that it has clinical significance. Sensorimotor gating deficits are widely observed in neuropsychiatric conditions that include schizophrenia, bipolar disorder (in particular acutely manic stages), and Tourette’s syndrome (39, 40). These deficits are likely to underlie clinical problems in distractibility due to the impaired ability to screen out irrelevant cues, cognitive fragmentation, and disintegrated thought (41, 42). “Dimensional” approaches to address brain function for a specific behavioral trait (e.g., sensorimotor gating or PPI in the present study) encompasses a much more efficient and effective strategy for discovery of drug targets for translation. Thus, the National Institute of Health states that such a dimensional approach, including the Research Domain Criteria (RDoC), may be a critical element in psychiatry in the overall scope of “Precision Medicine” (43).

In the present study, we used lithium as a pharmacological probe to address a key mechanism linking Bdnf transport and sensorimotor gating deficits in *Disc1*-LI mice. The beneficial action of lithium on the PPI deficits has been reported in more than one rodent model from multiple groups (44, 45). However, depending on the method of modeling, the effects of lithium are different. For example, chronic lithium treatment prevented amphetamine-induced PPI disruption, but not ketamine-induced PPI disruption (46). Addressing the context-dependent efficacy of lithium on PPI will be a future research question. Likewise, although the efficacy of lithium in treatment of patients with bipolar disorder has been well known and was also published for patients with HD as case reports (47, 48), its precise mechanisms regarding molecular targets and circuit specificity still remain elusive (49). In addition to currently accepted putative targets of lithium, such as inositol monophosphatase and glycogen synthase kinase-3 (GSK3) (50), further studies are awaited at the mechanistic levels, taking into account the cellular and circuit-wide functions of this compound. In this regard, the present study would provide important information for future translational efforts.

Here we focused on Bdnf transport as a plausible mechanism underlying sensorimotor gating. However, Disc1 also has more general roles in intracellular trafficking as a key component of the kinesin-1-driven axonal transport machinery, as *in vitro* studies have shown that it regulates transport of NUDEL/LIS1/14-3-3ε (28). Because Disc1 and Htt are rather ubiquitously expressed in the brain, one may also wonder why and how these proteins play a particularly important role in a specific behavior (*e.g*., sensorimotor gating or PPI) via regulation of Bdnf transport. We interpret this specificity because of the unique dependence of the striatum on the supply of Bdnf from the cortex. This uniqueness may emphasize the particular significance of this mechanism in the corticostriatal (PFC→DMS) projection and behavior.

We have recently reported pathological interaction of Disc1 and Htt in mood-associated symptoms in HD (15): we showed a “gain-of-function” of mutant Htt aberrantly sequestrated Disc1, but not phosphodiesterase 4 (Pde4), resulting in altered enzymatic activity of Pde4 that is to be controlled by Disc1 in a physiological condition. In contrast, in the present study, we demonstrated that “loss-of-function” of Disc1/Htt interaction in the PFC→DMS projection leads to a distinct cellular deficit (*i.e*., Bdnf transport), thereby affecting a distinct behavioral dimension *(i.e*., sensorimotor gating). This not only exemplifies multi-functional nature of both Disc1 and Htt, but also emphasizes a need to address each behavioral dimension based on specific molecular, cellular and circuit-wide mechanisms, highlighting the validity of dimensional approach in studying mechanisms underlying neuropsychiatric disorders.

Sensorimotor gating or PPI may be mediated by circuitries other than the cortico-striatal projection in which other molecular mediators possibly participate in other contexts. Our hypothesis-driven approach does not exclude this possibility. Elucidating all mechanisms for PPI may be too ambitious, which is beyond the scope of the present study. Nevertheless, the central mechanism that we present here at multiple levels in an integrated manner will provide an in-depth, fundamental insight into understanding sensorimotor gating in many brain disorders, such as schizophrenia and HD.

## Methods

### Mice

All animal studies were in accordance with guidelines for the care and use of laboratory animals issued by the Institutional Animal Care and Use Committees at Johns Hopkins, Duke, and City of Hope, as well as by the European Community (86/609/EEC) and the French National Committee (87/848). Generation of the *Disc1* LI mouse model is as described (19). Mice were group-housed in wire-topped clear plastic cages under a controlled environment with free access to food and water.

### *In vivo* electrophysiology

#### Electrode implantation surgery

Three- to four-month old mice were anesthetized with ketamine (100 mg/kg) and xylazine (10 mg/kg), placed in a stereotaxic device, and metal ground screws were secured to the skull above the cerebellum and anterior cranium. A total of 32 tungsten microwires were arranged in bundle arrays of 8–24 wires (each wire separated by at least 250 μm), and implanted using the following coordinates from Bregma: AP +2.6 mm, ML ±0.15 mm, DV −0.8 mm for PrL; AP +1.0 mm, ML ±0.9 mm, DV −2.2 mm for DSM. Full details of the procedures for electrode construction and surgical implantation have been previously described (51).

For optogenetic experiments, a Mono Fiberoptic Cannula coupled to a 2.5 mm metal ferrule (NA 0.22; 100 μm [inner diameter], 125 μm buffer [outer diameter], MFC_105/125-0.22, Doric Lenses, Quebec) was built directly into the PrL bundle. The tip of the fiber was secured 500 μm above the tip of the tungsten microwires. Implanted electrodes were anchored to ground screws using dental acrylic.

#### In vivo light stimulation

Laser output was controlled by digital input from the Cerebus acquisition system using custom written MATLAB scripts, and each light pulse was triggered by a 10 ms TTL signal. Experiments were performed with a laser power corresponding with 150 mw/mm^2^. Laser output was measured using a Power meter (Thorlabs, PM100D).

#### Prepulse inhibition (PPI) task

Startle responses were measured using a startle response (SR) chamber (medium animal enclosure; San Diego Instruments, CA). All mice were implanted with recording electrodes prior to experiments. On the experimental day, animals were anesthetized under isoflurane, and connected to the neurophysiological recording system. Following a 30 min recovery, mice were placed in the SR chamber, and neurophysiological and behavioral data were acquired as mice were subjected to the task. A 65 dB white noise stimulus was presented as background throughout the entire task. Five minutes following the beginning of the task, eighty-two auditory pulses (n = 30, 120 dB–40 ms stimulus [Startle Stimulus]; n = 30, 78 dB–40 ms stimulus followed by a 120 dB–40 ms stimulus 120 ms after the initial signal onset [PPI Stimulus]; n = 22, 78 dB-40 ms stimulus) were presented in pseudorandom order, with a pseudorandomized inter-pulse-interval ranging between 8 and 23s. Data recorded during the 78 dB-40 ms alone trials (n = 22) were not analyzed as part of this study since this stimulus was identical to the initial 120 ms of the PPI stimulus. Auditory stimuli were generated and delivered using custom written MATLAB (version R2010b MathWorks) scripts. In order to obtain the precise time stamps for stimulus presentation, acoustic data within the recording chamber was sampled continuously using a Digital Sound Level Meter (33-2055, Radio Shack, TX) and stored in real-time using our neurophysiological recording system. Startle responses were also recorded in real-time and stored along with our neurophysiological data.

#### Neurophysiological data acquisition

Neurophysiological recordings were performed during the PPI test. Neuronal activity was sampled at 30kHz, highpass filtered at 250Hz, sorted online, and stored using the Cerebus acquisition system (Blackrock Microsystems Inc., UT). Local field potential data was band pass filtered at 0.5-250Hz and stored at 1000Hz. Neuronal data were referenced online against a wire within the same brain area that did not exhibit an isolated neuron. At the end of the recording, cells were sorted again using an offline sorting algorithm (Plexon Inc., TX) to confirm the quality of the recorded cells. All neurophysiological recordings were referenced to a ground wire connected to both ground screws. Notably, wires tested from the two screws were iso-electric demonstrating that ground loops were not introduced by this design.

#### Construction of perievent time histogram (PETH)

Neuronal activity referenced to the onset of the 120 dB pulse was averaged in 20 ms bin, shifted by 1ms, and averaged across the 30 trials to construct the perievent time histogram for the startle and prepulse trials. Activity was referenced to the onset of each light pulse and averaged across 60 trials to construct the perievent time histogram for optogenetic stimulation experiments. Distributions of the histogram from the [−5,000 ms, −1,000 ms] interval were considered baseline activity. We then determined which 20ms bins, slid in 1ms steps during an epoch spanning from the [0 ms, 100 ms] interval for auditory stimulus trials or the [0 ms, 150 ms] interval for the optogenetic experiments met the criteria for modulation by a given stimulus. A unit was considered to be modulated by a given stimulus if at least consecutive 20 bins had firing rate either larger than a threshold of 99% above baseline activity or smaller than a threshold of 94% below baseline activity. This approach has been previously described (52).

#### Population gating effect

The magnitude firing rate fold change (unit gating effect) for each unit was calculated as: 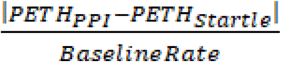; where the baseline rate for each neuron was given by the mean firing rate within the [−5,000 ms, −1,000 ms] interval prior to all startle and prepulse trials. Data was calculated for each 1ms bin of the PETH. For each temporal offset within the [−150 ms, 150 ms] interval, the population gating effect was given by the mean of the 50% of units, which exhibited the strongest gating effect at each offset. We employed a bootstrapping procedure in order to determine the significance threshold for each offset. The population gating effect within the [−5,000 ms, −1,000 ms] interval using the same units, which exhibited the strongest gating effects at a given offset was considered baseline activity. This process yielded 4,000 baseline values for a given 1ms bin, and a value corresponding with the 99% baseline threshold.

Population gating was considered significant within the [−150 ms, 150 ms] interval when the 99% confidence interval of the population gating effect for at least 20 consecutive bins was larger than a threshold of 99% above baseline activity. Next, we applied a second approach to confirm significant population gating effects. Unit gating effects were averaged across the temporal offsets that exhibited significant gating using our initial approach ([−114 ms, −95 ms]_PrL_, [−84 ms, −40 ms]_PrL_, and [−8 ms, 30 ms]_DMS_), and the 50% of units which exhibited the highest gating effects for a given interval were selected. Pair-wise comparisons were then performed between the mean gating effect of each unit within the significant intervals and the average gating effect observed for within the [−5,000 ms, −1,000 ms] interval across the unit population using a Wilcoxon signed-rank test at an alpha level of 0.05, followed by a Bonferroni correction for the three significant gating intervals. All three of these comparisons yielded statistically significant results.

#### Receiver operating characteristic (ROC) analysis to confirm neuronal gating deficit in Disc1 LI mice

To further quantify differences in neuronal gating signals across genotypes, we used a measurement based on receiver operator characteristic (ROC) analysis. ROC analysis is a widely utilized approach for classifying a neurons response to different stimuli (22). In contrast to the analysis applied to the startle and PPI PETH’s described above, ROC analysis provides an estimate of how the activity of a neuron differs between two stimuli on a trial by trial basis. The area under the ROC curve provides a measure of the extent to which a neuron discriminates two signals, where 0.5 corresponds to two overlapping distributions (no discrimination), and 0 or 1 corresponds to perfect discrimination (22).

### Histology

Mice were perfused using 1% PBS followed by 10% buffered formalin. After perfusion, the brain was carefully removed from the skull, washed with 10% buffered formalin and immersed in a 30% sucrose solution for 24-48 hours at 4°C. Brain slices (40 μm thick) were prepared using a cryostat, sections were stained with Nissl, mounted on slides using mounting medium, and covered with a coverslip.

### Ex vivo MRI

High resolution T2-weighted MRI of postmortem specimens was performed on an 11.7 Tesla vertical bore NMR spectrometer (Bruker Biospec, Inc.) by using a 15-mm diameter volume coil as the radiofrequency transmitter and receiver. Three-dimensional (3D) rapid acquisition with refocused echoes (RARE) sequence was used with the following parameters: TE 45 ms, TR 1500 ms, RARE factor 8; and 4 signal averages. The imaging field of view and matrix size were 16.0 × 12.0 × 10.0 and 200 × 152 × 128 (mm), respectively, and the native resolution was approximately 80 × 80 × 80 mm^3^. Total imaging time was 4 hours for each specimen.

### Behavioral testing

Behavioral testing was performed on male mice starting at 3–4 months of age with 1 week between the tests in the following order: open field, rotarod, and prepulse inhibition. We followed the same methods as in our previous publication (12), except for the rotarod test, which consisted of 1 day of training at 4 rpm for 5 min, followed by 3 consecutive days of testing at 4 to 40 rpm acceleration over a 5 min period per trial × 3 trials per day, with at least 30 min recovery between trials. WT, n ≥ 7; *Disc1* LI, n ≥ 8.

### Optogenetics

Three- to four-month-old mice were anesthetized with ketamine (100 mg/kg) and xylazine (10mg/kg), placed in a stereotaxic device, and injected with 3 μL virus in the PrL (AP +2.6 mm, ML±0.25 mm, DV −0.8 mm) using a 10 μl Hamilton micro syringe. Seven WT and 5 *Disc1* LI mice were unilaterally injected with pAAV-CaMKIIa-hChR2(H134R)-mCherry (CaMKII-Chr2) virus. Recording electrodes and stimulating fibers were surgically implanted 2 weeks following viral surgeries. Viral expression was confirmed electrophysiologically prior to neurophysiological recordings. Mice that failed to demonstrate neurophysiological responses to light stimulation (2/7 WT and 1/5 *Disc1* LI mice) were excluded from optogenetic studies.

### Bdnf

The amount of Bdnf in brain homogenates was measured by the sandwich ELISA method (BDNF Emax Immunoassay System, Promega). In brief, monoclonal BDNF antibody precoated on microtiter plates was used as a capturing antibody, and the captured Bdnf was quantified using chick anti-BDNF antibody and a chromogenic substrate, followed by detection with a plate reader at 450 nm wave-length, according to the manufacturer’s instructions. WT, n ≥ 8; *Disc1* LI, n ≥ 8. The levels of *Bdnf* mRNA at about 3 months of age were measured with TaqMan probe normalized to actin according to the published protocol (53). WT, n = 4; *Disc1* LI, n = 4.

### Bdnf transport assay with primary neuron culture

Mouse cortical neurons were prepared from E14.5 embryos and the Bdnf-mCherry expression construct was transfected 2 days prior to imaging using Lipofectamine 2000 (Invitrogen), according to the published protocol (27). Lithium (2 mM) was added to the culture medium 30 min before imaging. Time-lapse images were acquired every 100–200 ms up to 10 min, by using an IX81 fluorescence microscope (Olympus) equipped with a temperature-controlled unit, a gas-chamber (5% CO_2_, 37°C), a cooled charge-coupled device camera, and a 100x objective lens (NA = 1.3). Bdnf-mCherry-positive puncta were analyzed by Metamorph software (Molecular Devices). Three independent experiments were done; each time a pair of WT and *Disc1* LI mice within a litter were dissected to prepare primary cultures, and > 10 transfected cells were imaged for each genotype.

### Stereotaxic injection of AAV-Bdnf

Adeno-associated virus (AAV) was bilaterally injected into two sites the dorsomedial striatal locations: Site 1, AP: +0.9; ML +/− 1.2; DV −2.4 and −2.9. Site 2, AP: +0.4; ML +/− 1.2; DV −2.5 and −3.0. The stereotaxic injection was performed at 10 weeks of age with AAV expressing Bdnf or mCherry as control. Six weeks later, once the mice had recovered and Bdnf was expressed, the mice were tested in the PPI paradigm. WT + control, n = 11; *Disc1* LI + control, n = 10-14; *Disc1*-LI + Bdnf, n = 5-9 (after removal of negative PPIs).

### Chronic lithium treatment

Lithium was administered by daily i.p. injections (100 mg/kg) for 2 weeks. n = 5-8.

### Co-immunoprecipitation and Western blots

Triton X-100 (1%)-soluble brain extracts (~100 μg) were immunoprecipitated by anti-Htt (1 μg, mouse monoclonal, Millipore) and subsequently analyzed by Western blots using the antibodies indicated; anti-KIF5 (1:1,000, mouse monoclonal, Millipore), anti-p150^glued^ (1:1,000, mouse monoclonal, BD), anti-Disc1 (1:200, rabbit polyclonal, gift from Dr. Joseph Gogos, New York, USA), anti-HAP1 (1:200, mouse monoclonal, Millipore), and anti-Htt (1:2,000, mouse monoclonal, Millipore). Additional antibodies used were: phospho-Htt Ser-421 (1:1,000, rabbit polyclonal), AKT1 (1:1,000, rabbit monoclonal, Cell Signaling Technologies [CST]), phospho-AKT1 (1:200, rabbit monoclonal, CST).

### Statistics

The statistical test used is specified for each result. *p < 0.05, **p < 0.01, ***p < 0.001.

## Supporting information

## Author Contributions

Overall supervision of the entire group that includes multiple research teams: A.S. Conceptualization: H.J-P, K.Dz, T.T, A.S. Design of experiments: H.J-P, S.K, A.N.E, M.D.H, F.S, K.Dz. T.T. Performance of experiments: H.J-P, S.K., D.H, S-H.K, S.Z, R.T, Y.H-T, D.Z, A.S, A.G, J.B, B.M.K, Y.N-M, E.H, B.H, R.F, S.N, Z.H, T.H, W.C.W, K.I, K.Dz. Writing the manuscript: A.S, H.J-P, K.Dz, T.T. Funding directly for the experiments: A.S, K.Dz. Methodology: K.De, S.M. Intellectual contribution: N.J.B, M.T, M.D.H.

## Acknowledgments

We thank Drs. Francis Lee, Deqiang Jing, and Ravi Tharakan for discussion. We thank Drs. Melissa Landek-Salgado, Pamela Talalay and Ann West for critical reading and Ms. Yukiko Lema for organizing the manuscript and figures. We also thank Freeman Hrabowski, Robert and Jane Meyerhoff, and the Meyerhoff Scholarship Program (K.Dz.).

