## Supplementary material for "The cortico-striatal circuit regulates sensorimotor gating via Disc1/Huntingtin-mediated Bdnf transport"

### Supplemental Figure 1

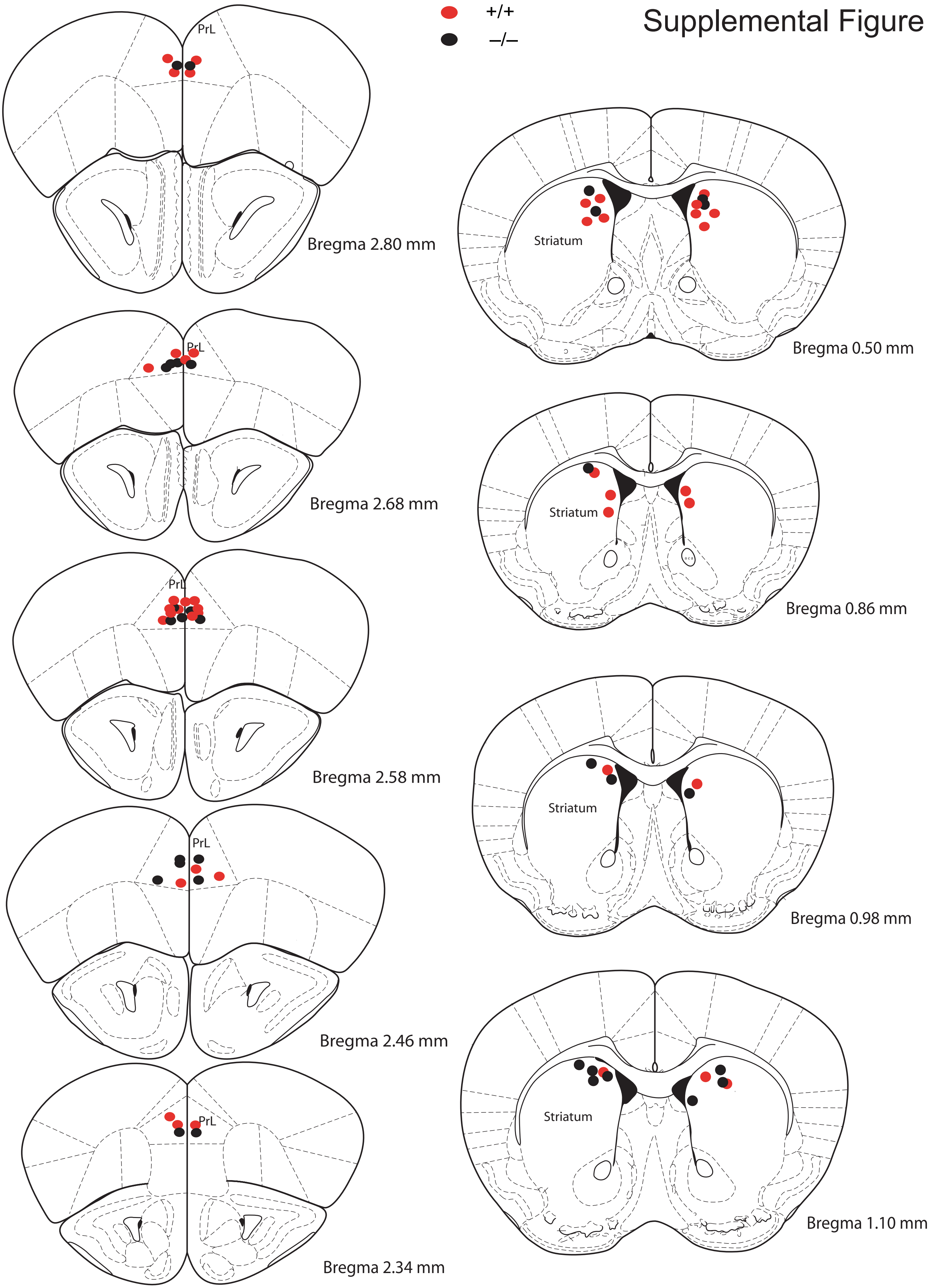

**Fig. S1. Verification of implantation sites.**
Mice were perfused following the completion of neurophysiological and behavioral studies. Brains were sliced on a cryostat, and stained for Nissl. Representative microwire implantation sites are shown at the corresponding AP coordinates. The right hemisphere was only targeted in animals implanted bilaterally. Atlas adapted from Paxinos & Franklin (Paxinos and Franklin, 2001).

A

| Volume ($mm^3$) | WT | | *Disc1* LI | | Comparison | |
|---|---|---|---|---|---|---|
| | **Average** | **Standard error** | **Average** | **Standard error** | ***Disc1* LI / WT** | **P** |
| Whole brain | 454.94 | 11.36 | 424.84 | 6.91 | 0.93 | 0.08 |
| **Cerebellum** | 59.30 | 1.43 | 52.74 | 1.49 | 0.89 | **0.02** |
| Hippocampus | 22.28 | 1.03 | 21.21 | 0.75 | 0.95 | 0.46 |
| Hypothalamus | 10.24 | 0.40 | 10.24 | 0.48 | 1.00 | 1.00 |
| Neocortex | 95.95 | 5.12 | 89.87 | 1.35 | 0.94 | 0.35 |
| **Striatum** | 20.63 | 0.42 | 18.90 | 0.39 | 0.92 | **0.02** |
| Thalamus | 21.55 | 0.61 | 19.61 | 1.29 | 0.91 | 0.23 |
| Ventricles | 5.57 | 0.39 | 5.72 | 0.29 | 1.03 | 0.78 |

B

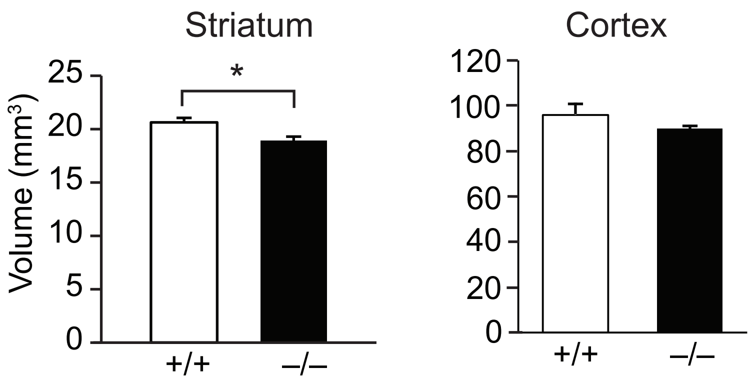


**Fig. S2. Gross brain anatomy by structural magnetic resonance imaging.**
(A) Volumes of the whole brain and regions of interest (average and standard error) as determined by structural MRI at 6 months of age.
The volumes were compared by dividing the average *Disc1* LI volume by the average WT volume and by student's t-test. The P<0.05 results are bolded.
(B) The striatum of *Disc1* LI **(-/-)** mice was significantly smaller in volume compared to that in WT (+/+) littermates (left). There were no significant differences in the volume of the cortex (right). *P<0.05 using student's t-test.

Supplemental Figure 3

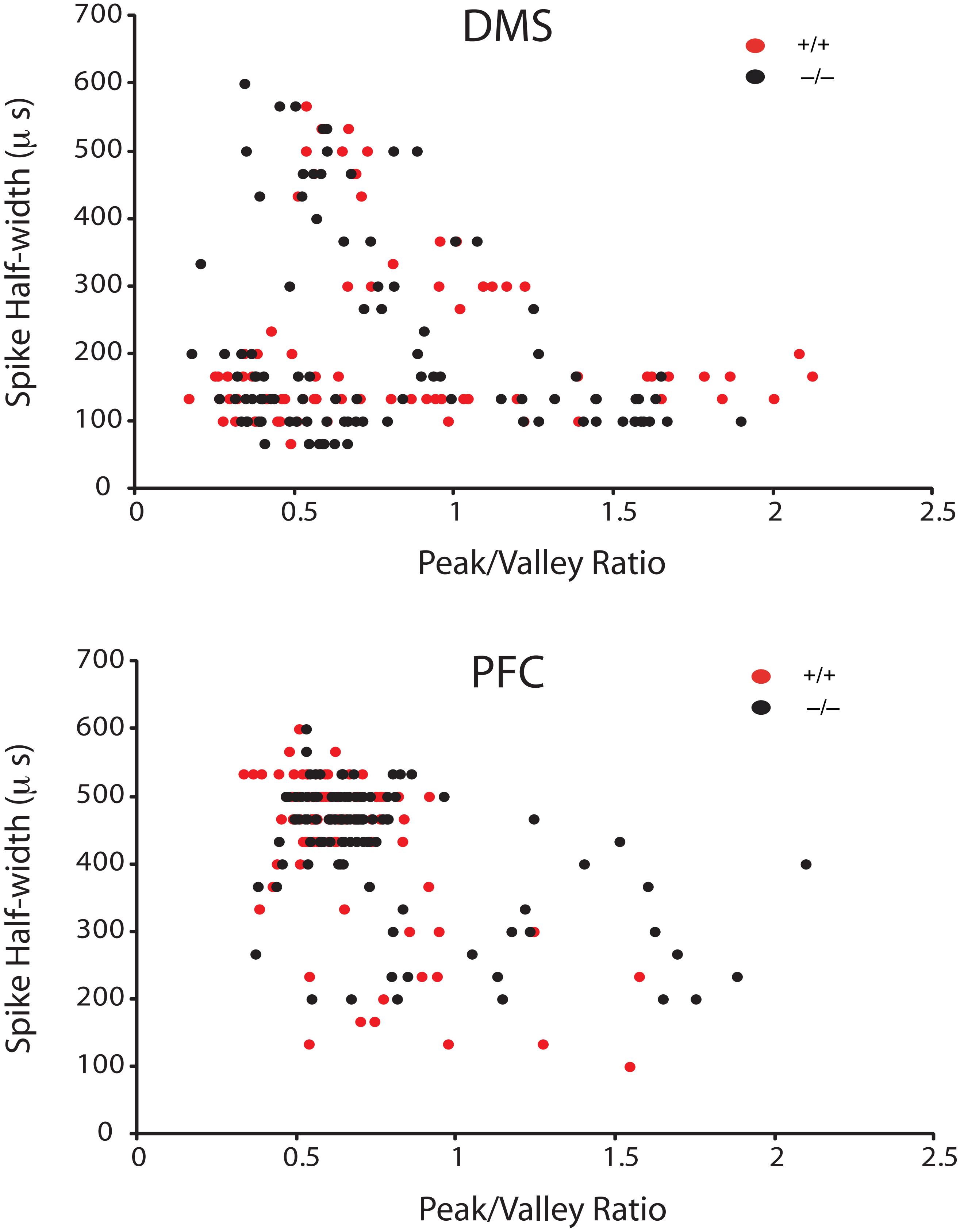


**Fig. S3. Waveform properties of DMS and PFC cells recorded in Disc1 LI and WT mice.** Note the overlap between teh distributions from Disc1 LI (-/-) and WT (+/+) littermate controls.

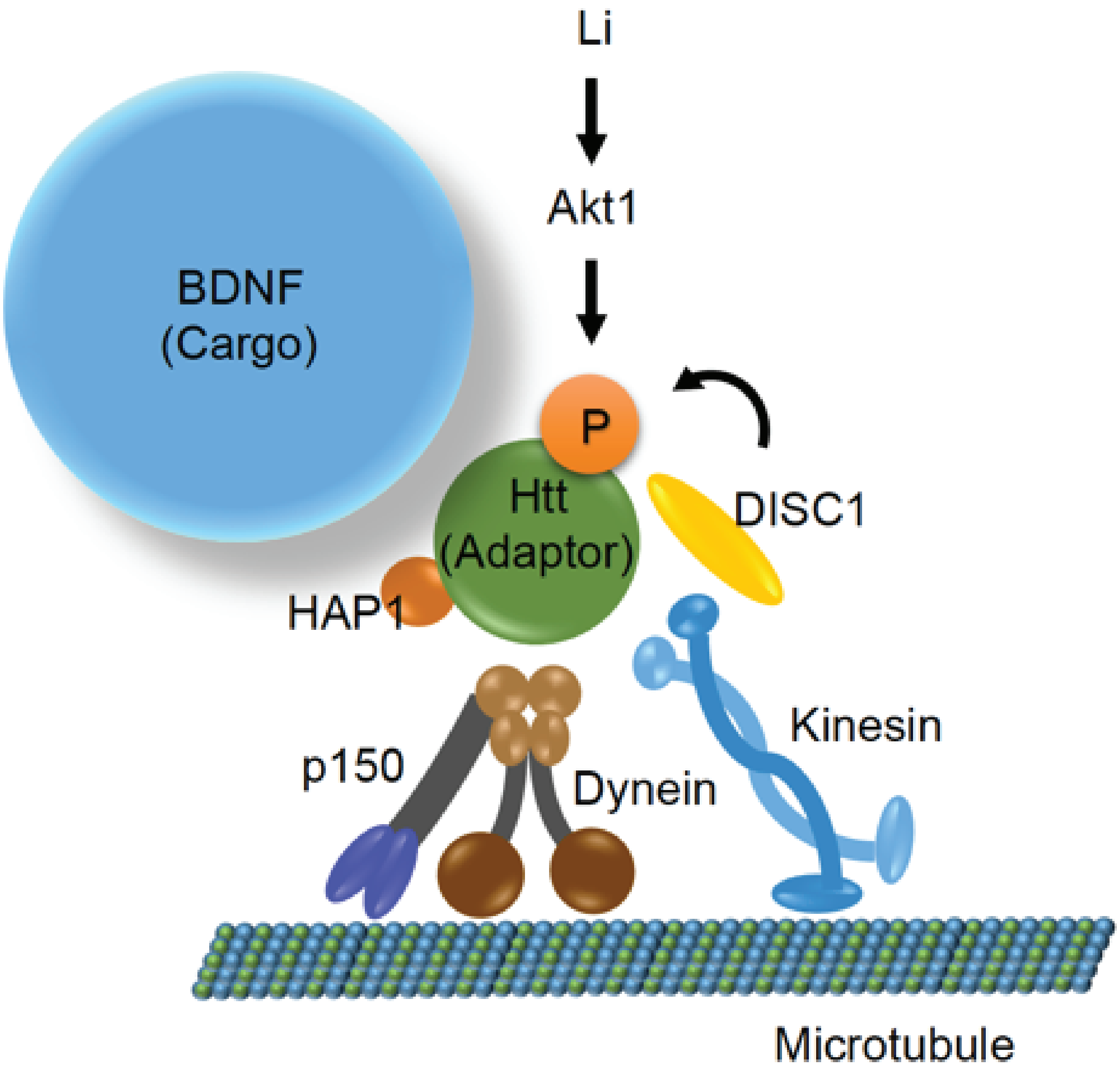


**Fig. S4. Schematic model for Disc1 and Li in supporting Htt-mediated Bdnf transport.** Bdnf-containing cargo is linked to the motor machinery (kinesin and dynactin) via Htt adaptor protein and thereby transported along the cortico-striatal tract. Disc1 supports Bdnf transport by facilitating the complex formation among Htt, cargo (Bdnf) and motors, in part through augmentation of Ser-421 phosphorylation of Htt. Lithium (Li) could also enhance this complex formation via Ser-421 phosphorylation of Htt, possibly through upregulation of Akt1 activity or Akt1 recruitment.

Supplemental Figure 5

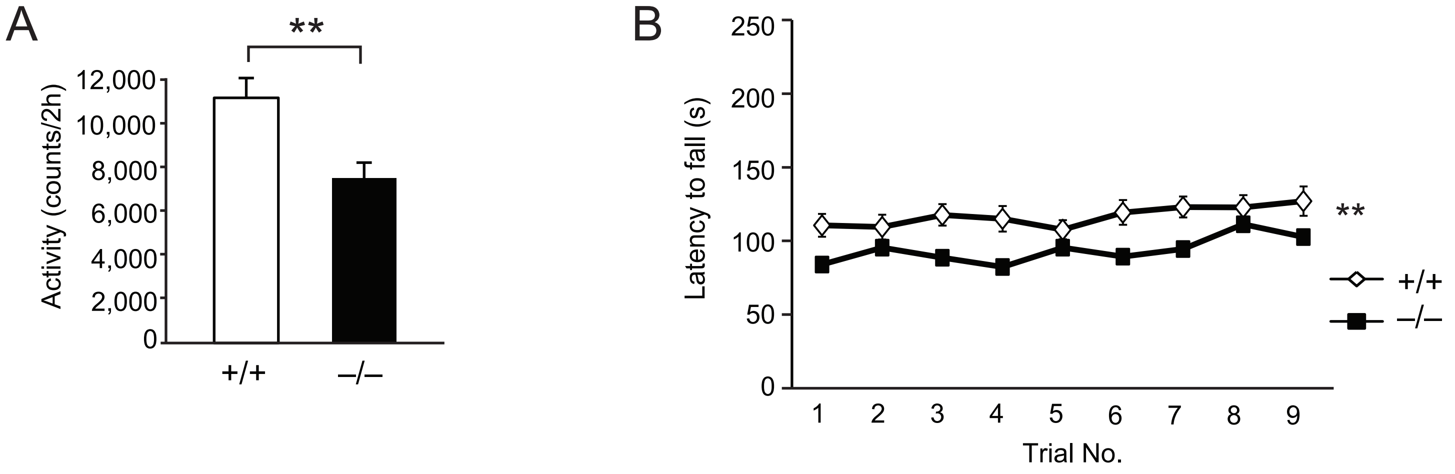


**Fig. S5. Additional behavioral deficits in Disc1 LI mice compared to WT littermates.**
(A) *Disc1* LI mice (-/-) displayed a significant reduction in overall locomotor activity in the open field compared to WT littermates (+/+). **P<0.01 using student's t-test.
(B) *Disc1* LI mice (-/-) displayed impaired motor function compared to WT littermates (+/+) as measured by the rotarod test. **P<0.01 using repeated measures ANOVA.
